# RgCop-A regularized copula based method for gene selection in single cell rna-seq data

**DOI:** 10.1101/2020.12.23.424205

**Authors:** Snehalika Lall, Sumanta Ray, Sanghamitra Bandyopadhyay

## Abstract

Gene selection in unannotated large single cell RNA sequencing (scRNA-seq) data is important and crucial step in the preliminary step of downstream analysis. The existing approaches are primarily based on high variation (highly variable genes) or significant high expression (highly expressed genes) failed to provide stable and predictive feature set due to technical noise present in the data. Here, we propose *RgCop*, a novel **r**e**g**ularized **cop**ula based method for gene selection from large single cell RNA-seq data. *RgCop* utilizes copula correlation (*Ccor*), a robust equitable dependence measure that captures multivariate dependency among a set of genes in single cell expression data. We raise an objective function by adding a *l*_1_ regularization term with *Ccor* to penalizes the redundant co-efficient of features/genes, resulting non-redundant effective features/genes set. Results show a significant improvement in the clustering/classification performance of real life scRNA-seq data over the other state-of-the-art. *RgCop* performs extremely well in capturing dependence among the features of noisy data due to the scale invariant property of copula, thereby improving the stability of the method. Moreover, the differentially expressed (DE) genes identified from the clusters of scRNA-seq data are found to provide an accurate annotation of cells. Finally, the features/genes obtained from *RgCop* can able to annotate the unknown cells with high accuracy.

**Availability:** Corresponding software is available in: https://github.com/Snehalikalall/RgCop

## 1 Introduction

With the advancement of single cell RNA-seq (scRNA-seq) technology a wealth of data has been generated allowing researchers to measure and quantify RNA levels on large scales [1]. This is important to get valuable insights regarding the complex properties of cell type, which is required for understanding the cell development and disease. A key goal of single cell RNA-seq analysis is to annotate cells within the cell clusters as efficiently as possible. To do this, the basic goal would be to select a few informative genes that can lead to a pure and homogeneous clustering of cells [2, 3]. The task of selecting effective genes among all gene panel that can precisely discriminate cell type labels can be regarded as a combinatorially hard problem.

The usual approach for annotating cells is to cluster them into different groups which are further annotated to determine the identity of cells [4, 5]. This is considered a popular and unsupervised way of annotating different types of cells present in a large population of scRNA-seq data [6–8]. The general pipeline of downstream analysis of scRNA-seq data typically goes through several steps. Starting from the processing of the raw count matrix, the scRNA-seq data is going through the following steps: i) normalization (and quality control) of the count matrix ii) feature selection, and iii) dimensionality reduction [2, 9]. The first step is necessary to adjust discrepancies between samples of individual cells. Several quality measures are also applied to reduces the skewness of the data. The next step identifies the most relevant features/genes from the normalized data. The relevant genes are either selected by identifying the variation (highly variable genes) [10] or can be selected by calculating the expression levels across all cells which are higher than the average value (highly expressed genes).

The selection of top genes has a good impact on the cell clustering process in the later stage of downstream analysis. A good clustering can be ensured by the following characteristics of features/genes: the features/genes should have useful information about the biology of the system, while not including features containing any random noise. Thereby the selected features/genes reduce the data size while preserving the useful biological structure, reducing the computational cost of later steps.

The usual approach of gene selection is based on the high variability of the gene expression label of scRNA-seq data. This process is simple and suffers from several disadvantages: i) as the expression variability is dependent on pseudo-count, it can introduce biases in the data, ii) Next, PCA is applied in downstream analysis for dimensionality reduction which is not suitable for sparse and skewed scRNA-seq data.

In this paper, we present a method for finding the most informative features/genes from large scRNA-seq datasets based on a robust and equitable dependence measure called copula correlation (Ccor). Although the major applications of copula can be found in the domain of time series, finance, and economics, but it is now ripe for application in different domain of bioinformatics such as for modeling directional dependence of genes [11], finding differentially co-expressed genes [12] high dimensional clustering for identification of sub-populations of patients selection [13] and many more. However, the application of copula in single cell domain, particularly on gene selection is less explored.

In this paper, we show that employing a simple *l*_1_ regularization term with the proposed objective function, will improve the performance of any clustering/classification model significantly. The objective function is utilized a new robust-equitable copula correlation (Ccor) measure on one hand and a regularization term to control the coefficient on other hand. Thus it is robust due to regularization and not susceptible to noise due to scale-invariant property copula. *RgCop* has major advantages both in clustering/classifying unknown samples and in the identification of meaningful marker detection. The latter point is addressed because novel marker genes for different cell types are identified with the cell clusters. Biologically meaningful marker selection is usually an important step in the downstream analysis of scRNA-seq analysis. This depends on the homogeneity of the cell clusters identified after the gene selection stage. Our proposed method selects the most informative genes that ultimately leads to a homogeneous grouping of cells of the large scRNA-seq data.

Beyond selecting a good informative gene set that leads good clustering/classification of cells, we also demonstrate that our method performs well in completely independent data of the same tissue. We demonstrate this by evaluating the performance of the selected features in completely unknown test samples. We observed that the selected features are equally effective for clustering/classifying the unknown test samples. We further carry out a simulation study on eight Gaussian Mixture datasets to establish the effectiveness of the proposed method. The results show that the proposed method not only select genes with high accuracy, but is also robust and less susceptible to noise.

### Summary of contributions

The main contributions of the paper are summarized below:

– We provide one of the first regularized copula correlation (Ccor) based robust gene selection approaches from large scRNA-seq expression data. This robustness is a characteristic of our proposed objective function. The method also works equally well for the small sample and large feature scRNA-seq data.
– We derive a new objective function using the copula correlation and regularization term and theoretically prove that the selected feature set is optimal with respect to the minimum redundancy criterion
– The objective function of *RgCop* is so designed that it can simultaneously maximize the relevancy criterion and minimize the redundancy criterion among the two sets of features/genes. The regularization term is also added with the objective function so as to control the large coefficient of the relevancy term.
– *RgCop* can able to effectively cluster/classify unseen scRNA-seq expression data. Annotating unknown cells is crucial and is the final goal of scRNA-seq data analysis. We demonstrate the applicability of our framework in this case. We demonstrate that the selected features are effective for clustering completely unseen test data. The annotation of cells can also be done in a supervised way if one can train a classification model with the selected features.
– Our method is less sensitive (robust) against the noises present in the scRNA-seq data. Our objective function uses copula-correlation, a robust-equitable measure which has the advantage of capturing the multivariate dependency between two sets of random variables.

## Results

### Workflow

Figure 1 provides a workflow of the whole analysis performed here. Following subsections discussed the important steps:

**Fig 1.**
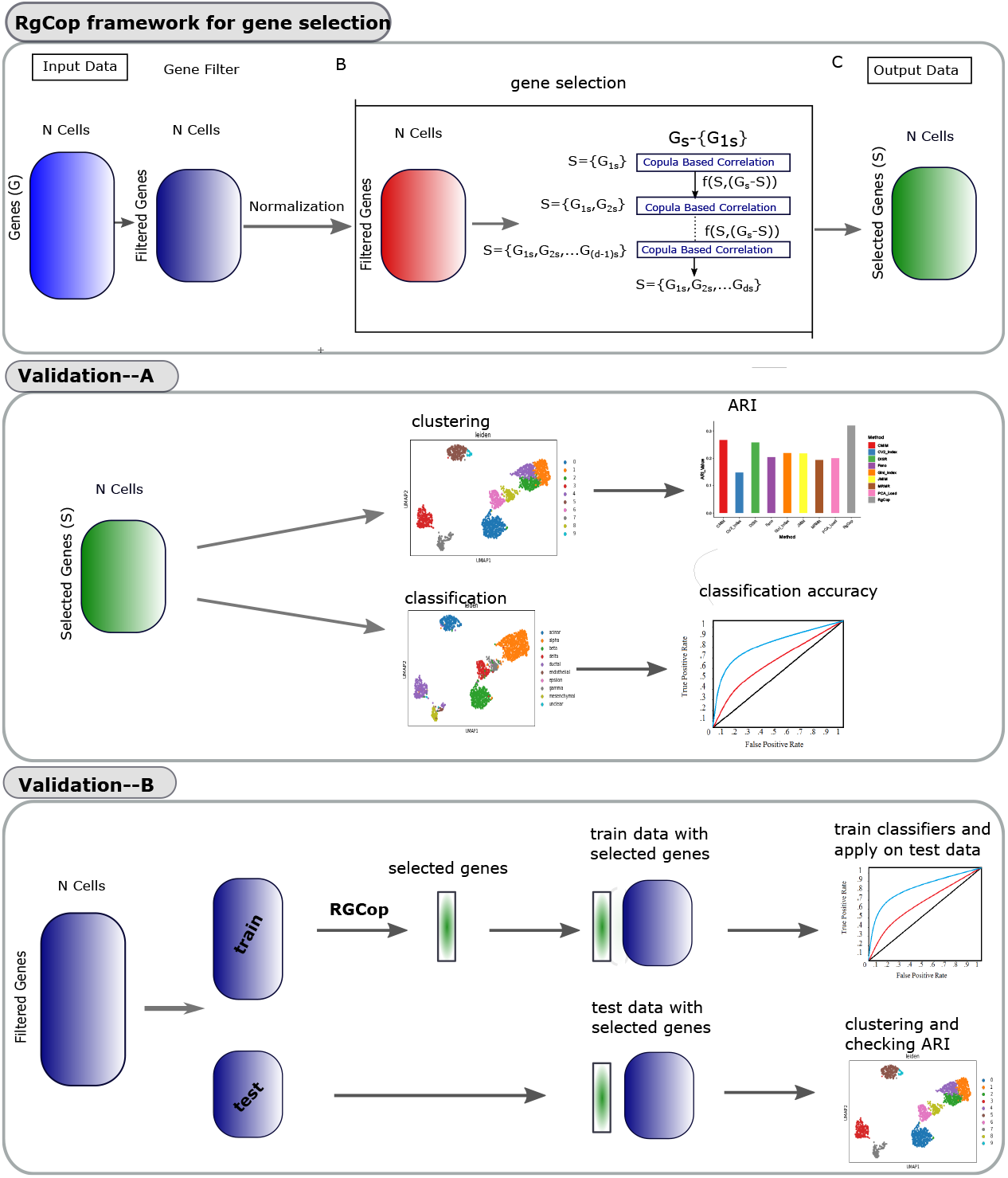
The whole workflow of the methodology: *RgCop* framework for gene selection is provided in the top panel. Clustering and classification is performed with the genes obtained from *RgCop* to validate the method (shown in middle panel). *RgCop* is validated for detection of unknown sample by splitting the data into train-test ratio of 7:3 (shown in the bottom panel). The test data is utilized for validation of the selected genes by *RgCop*.

#### A. Preprocessing of raw datasets

See -A of panel-‘*RgCop* framework for gene selection’ of figure 1. Raw scRNA-seq datasets are obtained from public data sources. The counts matrix 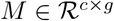, where *c* is number of cells and *g* represents the number of genes, is normalized using a transformation method (Linnorm) [14]. We choose that cells which have more than a thousand genes expression values (non zero values) and choose that genes which have the minimum read count greater than 5 in at least 10% among all the cells. *log*_2_ normalization is employed on the transformed matrix by adding one as a pseudo count.

#### B. RgCop framework for feature selection

See -B of panel-‘*RgCop* framework for gene selection’ of figure 1. The preprocessed data is used in the proposed copula-correlation (*Ccor*) based feature/gene selection models. First, a feature ranking is performed based on the *Ccor* scores between all features and class labels. We assume the feature having a larger *Ccor* value is the most relevant one and we include it in the selected list. Next, *Ccor* is computed between the selected relevant features and the remaining features. The feature with a minimum score is called the most essential (and not redundant) feature and included in the selected list. The process continued in an iterative way by including the most relevant and minimum redundant features in each step every time in the list. Feature selection in this way ensures the list of genes will be optimal (see proof of correctness). An *l*_1_ regularization term is added with the objective function to penalize the large coefficient of relevancy term. The resulting matrix with selected features is utlized for further doenstream analysis.

#### C. Validation through clustering

See panel-‘validation-A’ of the figure 1. We adopt the conventional clustering steps of scanpy [15] package to cluster the resulting matrix obtained from the previous step. We employed two clustering techniques (SC3 [4], and Leiden clustering [16]) for clustering the neighborhood graph of cells. To validate the clusters we utilize the Adjusted Rand Index (ARI) metric which is usually used as a measure of agreement between two partitions. We compare the ARI score of *RgCop* with different state-of-the-art unsupervised feature selection method.

#### D. Validation through classification

See panel-‘validation-A’ of the figure 1. We validate the selected features by employing several classifiers to train the resulting matrix obtained from step-B of the workflow. The features are selected by several supervised feature selection algorithm and the classification accuracy are compared with *RgCop*.

#### E. Annotating unknown cells

See panel-‘validation-B’ of the figure 1. For cells of the unknown type, *RgCop* can able to accurately cluster/classify the cells using the genes selected in the previous step. The filtered and preprocessed data is divided into train-test ratio 7:3 and the train set is utilized to obtain the selected features using *RgCop*. Several classifier models are trained on the train set with the selected feature set and applied to the test set. The test data with selected features are also used for clustering. This provides the validation of our approach to work in practice.

#### F. Marker identification

We detect highly differentially expressed (DE) genes within each cluster obtained from step-C in the workflow. Here we utilizing Wilcoxon Ranksum test to identify DE genes in each cluster. The top five DE genes are chosen from each cluster according to their p-values.

### Performance in synthetic Gaussian mixture data

For single cell clustering the most common challenge is to discriminating samples between major cell types and its sub-types. Samples of similar cell types tend to overlap within one cluster, discriminating of which required sophisticated method that can extract features from overlapped samples. To explore whether RgCop can address this issue we apply it on synthetic data with overlapping and nonoverlapping classes.

We generate four pair of synthetic Gaussian mixture datasets, each pair consisting of overlapping and non-overlapping classes (see supplementary for description of data generation).Figure 2, panel-A illustrates a 2D pictorial representation of the synthetic gaussian mixture datasets.

**Fig 2.**
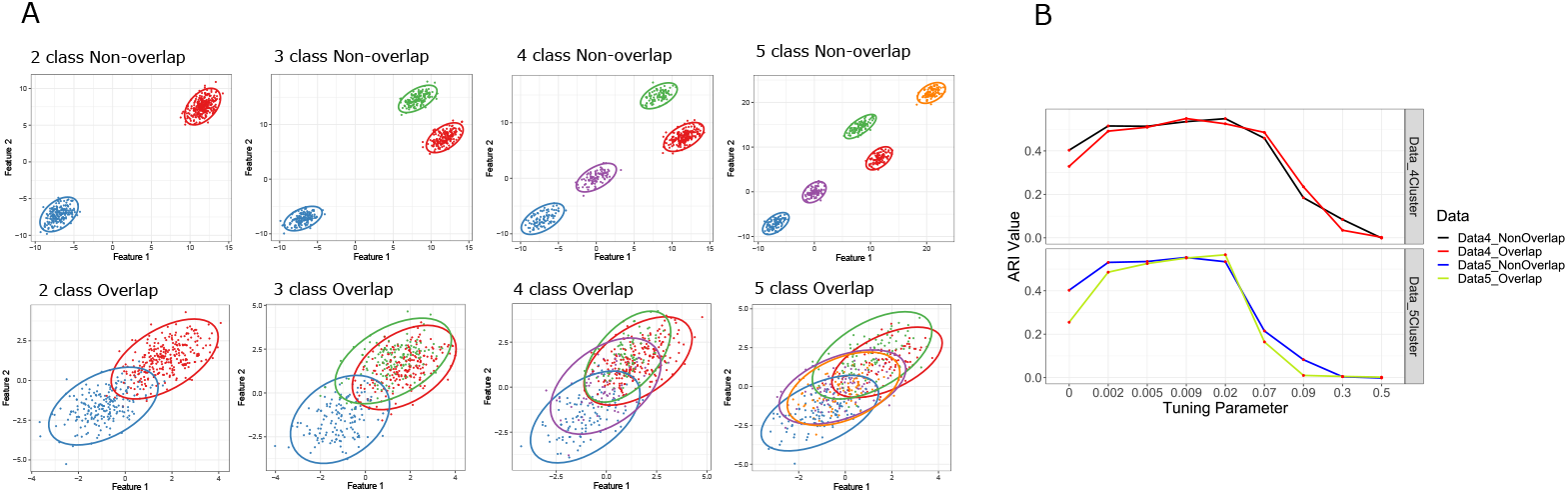
Figure shows tSNE visualization of eight synthetic Gaussian Mixture Datasets (Panel-A). The first one represents datasets of non-overlapping classes whereas the latter represents the same for overlapping classes. Panel-B shows the ARI scores of clustering results of synthetic data including four overlapped and non-overlapped clusters. *RgCop* is employed with 9 tuning parameters within the range [0,0.5]

We select top 100 features from the generated datasets. *k*-means clustering is employed to cluster the data with selected features. Clustering performance is evaluated using Adjusted Rand Index (ARI), a widely used metric to measure concordance between the predicted clusters and the known groups. Figure 2, panel-B describes the ARI values collected from the clustering results of the data with four and five overlapping and non-overlapping classes. The higher values of ARI suggests RgCop can perform well in data consiting of overlapping and non-overlapping classes.

To tune the regularization parameter *γ* (see equation 6), the clustering process is repeated for nine set of values ranging from 0 to 0.5 (*γ* = {0, 0.002, 0.005, 0.009, 0.02, 0.07, 0.09, 0.3, 0.5}). Clustering results is reported in figure 2, panel-B, which shows *RgCop* produces highest ARI value in the range *γ* ∈ [0.005, 0.02]. This values are utilized on the real datasets for gene selection.

### Comparison with State-of-the-art

We compared the efficacy of *RgCop* by comparing with four well known techniques for identifying highly dispersed genes in scRNA-seq data: *Gini Clust* [17], *PCA Loading* [18], *CV*^2^ *Index* and *Fano Factor* [19]. We also compared the performance of *RgCop* with four widely used supervised feature selection techniques:CMIM [20], JMIM [21], DISR [22], MRMR [23]. A short description of competing methods and parameter settings is described in the supplementary text.

#### Clustering performance on real dataset using unsupervised method

Here single cell Consensus clustering (SC3) method [4] is employed for clustering expression matrix with selected features. Figure 3, panel-A illustrates the boxplots of ARI Values of the clustering results on Yan, Muraro, and Pollen datasets. We vary the number of selected features in the range from 20 to 100 and compute the ARI scores for each method. It can be seen from the figure that *RgCop* achieves high ARI values in almost all the three datasets. For the Yan dataset, while the performance of other methods is relatively low, *RgCop* achieves a good ARI value, demonstrating the capability of *RgCop* to perform in small sample data. We also create a visualization of the clustering performance of *RgCop* in Muraro, yan, and Pollen datasets. Figure 3 panel- B, shows two dimensional t-SNE plot of predicted clusters and their original labels. Panel-C of this figure shows heatmaps of cell × cell consensus matrix representing how often a pair of cells is assigned to the same cluster considering the average of clustering results from all combinations of clustering parameter [4]. Zero score (blue) means two cells are always assigned to different clusters, while score ‘1’ (red) represents two cells are always within the same cluster. The clustering will be perfect if all diagonal blocks are completely red and off-diagonals are blue. A perfect match between the predicted clusters and the original labels can be seen from panel-B and panel-C of figure 3.

**Fig 3.**
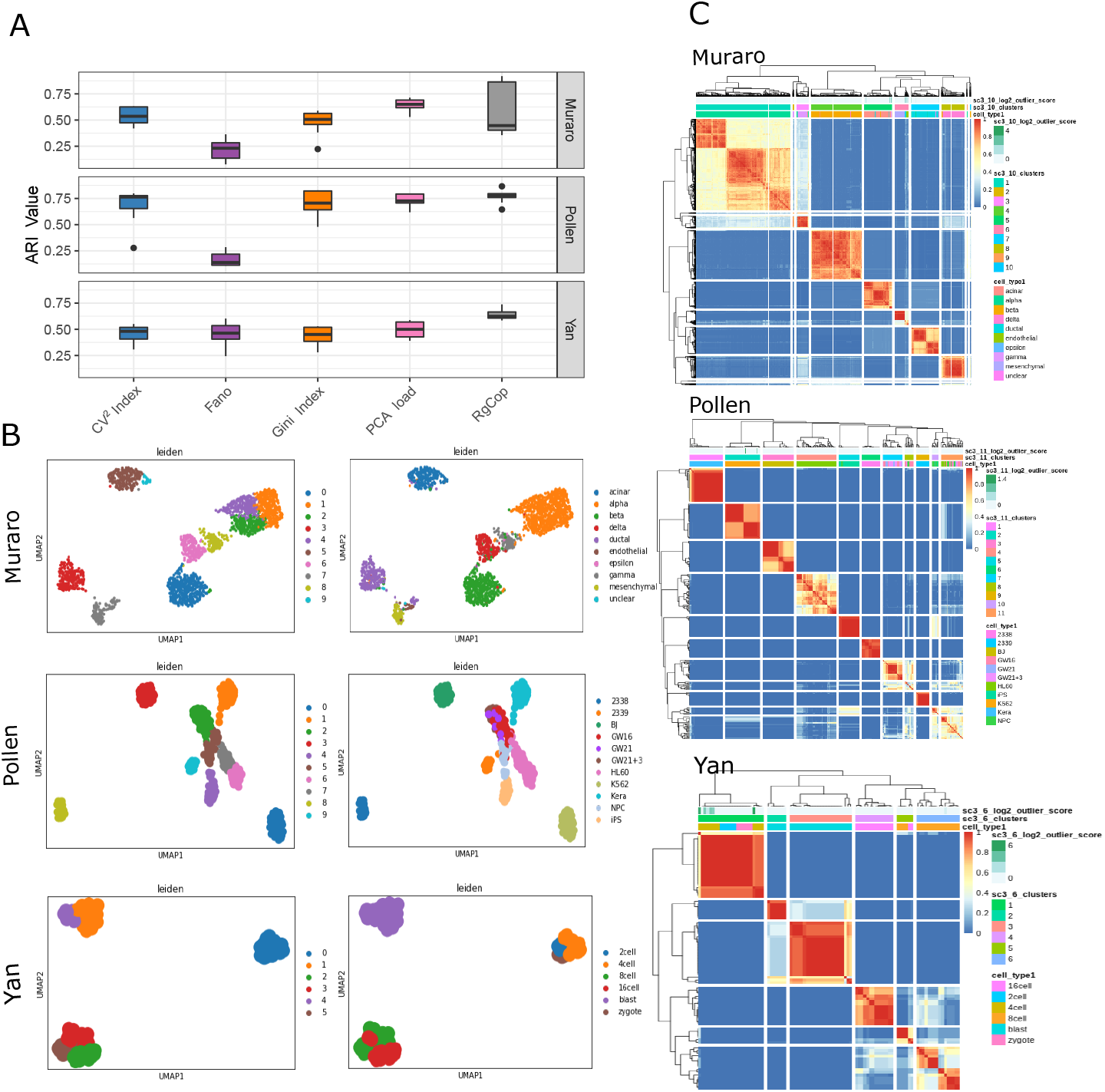
Figure shows the comparisons of clustering performance. Panel-A shows the boxplot of ARI values computed from the clustering results of each competing method. Each box represents ten ARI scores of clustering results for selected 10 sets of features ranging from 10 to 100. Panel-B shows the t-sne visualization of clustering results of three datasets for *RgCop*. Panel-C shows the consensus clustering plots of obtained clusters from *RgCop*.

#### Classification performance on real dataset using supervised method

We compare *RgCop* with four well known supervised feature selection methods and compute the classification accuracy. Four widely used classifiers are considered in our work, Support Vector Machine (*SVM*), Neural Network (*NN*), and Gradient Boosting Machine (*GBM*) for learning the expression matrix with selected features. Table 1 shows the average test accuracy and the corresponding standard errors over 50 runs for each of the competing methods.

**Table 1.** Classification results on Real Datasets using Supervised Methods

| Classifier | Dataset Name | MRMR | DISR | JMIM | CMIM | $RgCop$<br>($\gamma = 0.009$) |
| --- | --- | --- | --- | --- | --- | --- |
| 4*GBM | Muraro | 0.90 ± 0.05 | 0.89 ± 0.04 | 0.89 ± 0.010 | 0.85 ± 0.02 | **0.96 ± 0.009** |
| | Pollen | 0.75 ± 0.03 | 0.76 ± 0.01 | 0.75 ± 0.05 | 0.74 ± 0.02 | **0.88 ± 0.002** |
| | Yan | 1 ± 0 | 0.99 ± 0.02 | 0.98 ± 0.02 | **1 ± 0** | 0.97 ± 0.01 |
| 4*NNET | Muraro | 0.82 ± 0.02 | 0.81 ± 0.01 | 0.81 ± 0.05 | 0.84 ± 0.03 | **0.89 ± 0.07** |
| | Pollen | 0.71 ± 0.01 | 0.72 ± 0.003 | 0.69 ± 0.02 | 0.70 ± 0.01 | **0.85 ± 0.02** |
| | Yan | 0.99 ± 0.01 | **0.99 ± 0.02** | 0.99 ± 0.03 | 0.98 ± 0.003 | 0.98 ± 0.02 |
| 4*SVM | Muraro | 0.87 ± 0.04 | 0.88 ± 0.01 | 0.87 ± 0.011 | 0.89 ± 0.01 | **0.95 ± 0.01** |
| | Pollen | 0.74 ± 0.06 | 0.75 ± 0.03 | 0.70 ± 0.03 | 0.71 ± 0.01 | **0.87 ± 0.001** |
| | Yan | 1 ± 0 | 0.99 ± 0.02 | **1 ± 0** | 0.99 ± 0.04 | 0.98 ± 0.003 |

### Stability performance

Protocols for preprocessing of scRNA-seq data are complex and often suffer from technical biases that vary across cells. This causes biases in the downstream analysis if the noise is not properly handled. *RgCop* utilizes copula which is well known for its scale invariance property that makes it robust against noise in the data. To show the performance of *RgCop* in noisy data, white Gaussian noise with a mean (*μ*= 0) and standard deviation 1 is mixed to each gene/feature of a dataset. The function *Add.Gaussian.noise* of R package RMThreshold is used to generate Gaussian noise. Next, relevant 100 genes/features are chosen from each of the noisy datasets, and the percentage of matching is computed with the original genes/feature sets. We define a matching feature score (percentage) as follows *match_score* = ((*N – D*)*/N*) * 100, where *N* represents the total number of features, and *D* represents the number of feature discrepancies between the original and noisy dataset. We perform 10 trials, each contains 100 such experiments. For a competing method, each trial gives 100 scores for one dataset and the median of these scores are shown in figure 4. Each row of the figure shows bar plots of the median values for three scRNA-seq datasets. It can be observed from the figure that *RgCop* achieves better *match_score* for all the datasets, particularly for small sample data.

**Fig 4.**
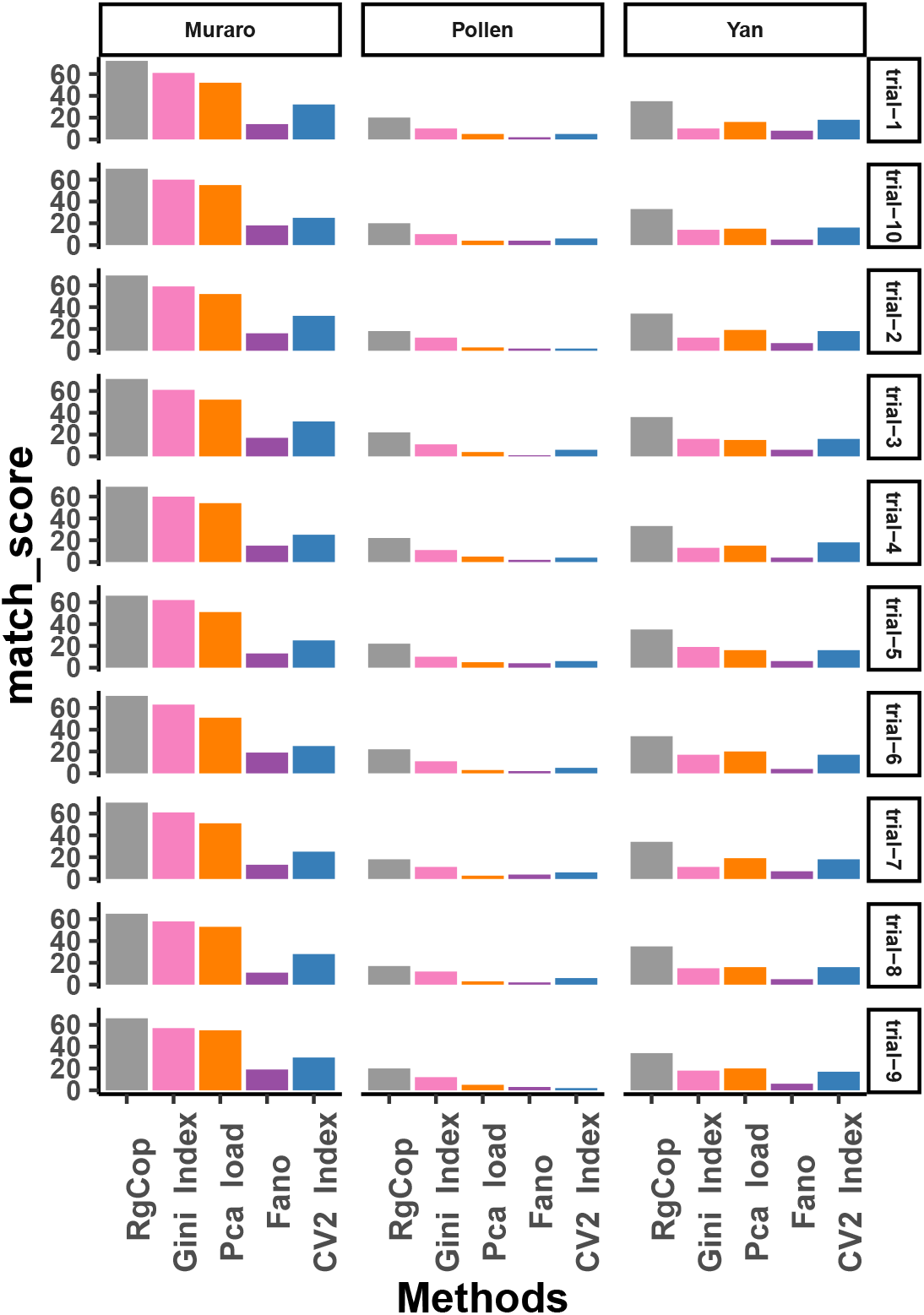
Figure shows the median of *match_score* (percentage) of five different competing methods. 10 trials are performed with 100 iteration in each trial to compute the median of *match_score*.

### Annotation of unknown samples

Predicting unknown sample is the final goal of any scRNA-seq analysis pipeline. Here, we addressed this problem by using a cross validation approach. We first split the data in training and test set in the ratio 7:3. Next, top 100 informative genes are chosen using *RgCop* from the training dataset. The clustering performance of *RgCop* is computed on the test data using the selected genes from the trained dataset. We repeat the procedure 50 times with a random split of train-test (7:3 ratio) for each data. The table 2 represents the mean and standard deviation of the ARI scores using *RgCop*. We also train three classifiers SVM, NN, and GBM on the train datasets with the selected feature. The aim is to see whether the trained model can predict the cell type of sample from the test data. Table 2 shows the classification accuracy of three classifiers.

**Table 2.**
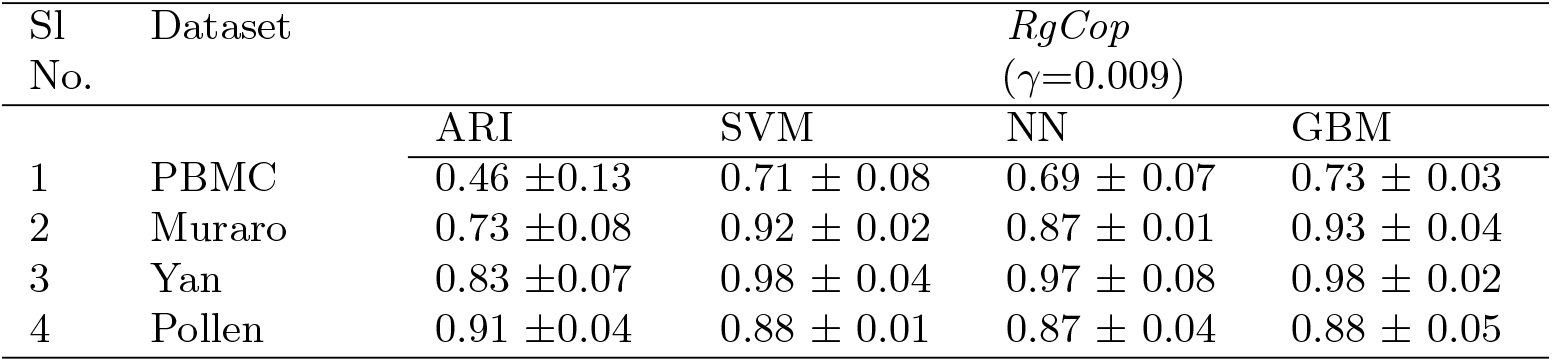
Adjusted Rand Index measured on the clustering results on unknown test samples

| Sl No. | Dataset | $RgCop$<br>($\gamma=0.009$) | | | |
| --- | --- | --- | --- | --- | --- |
|  |  | ARI | SVM | NN | GBM |
| 1 | PBMC | 0.46 ± 0.13 | 0.71 ± 0.08 | 0.69 ± 0.07 | 0.73 ± 0.03 |
| 2 | Muraro | 0.73 ± 0.08 | 0.92 ± 0.02 | 0.87 ± 0.01 | 0.93 ± 0.04 |
| 3 | Yan | 0.83 ± 0.07 | 0.98 ± 0.04 | 0.97 ± 0.08 | 0.98 ± 0.02 |
| 4 | Pollen | 0.91 ± 0.04 | 0.88 ± 0.01 | 0.87 ± 0.04 | 0.88 ± 0.05 |

### Marker gene selection

We have chosen marker genes (DE genes) for different cell types from the clustering results. Differentially Expressed (DE) genes are identified from every cluster using Wilcoxon rank-sum test. We use this to directly assess the separation between distributions of expression profiles of genes from different clusters. Figure 5 illustrates the top five DE genes from each cluster of Pollen dataset (panel-A), and Yan dataset (panel-B). The higher expression values of the top five DE genes (shown in the heatmap of panel-A, B) for a particular cluster suggests the presence of marker genes within the selected gene sets. The results are detectable from violin plots of the expression profiles of top DE genes within each cluster (figure 5 panel-A, and -B).

**Fig 5.**
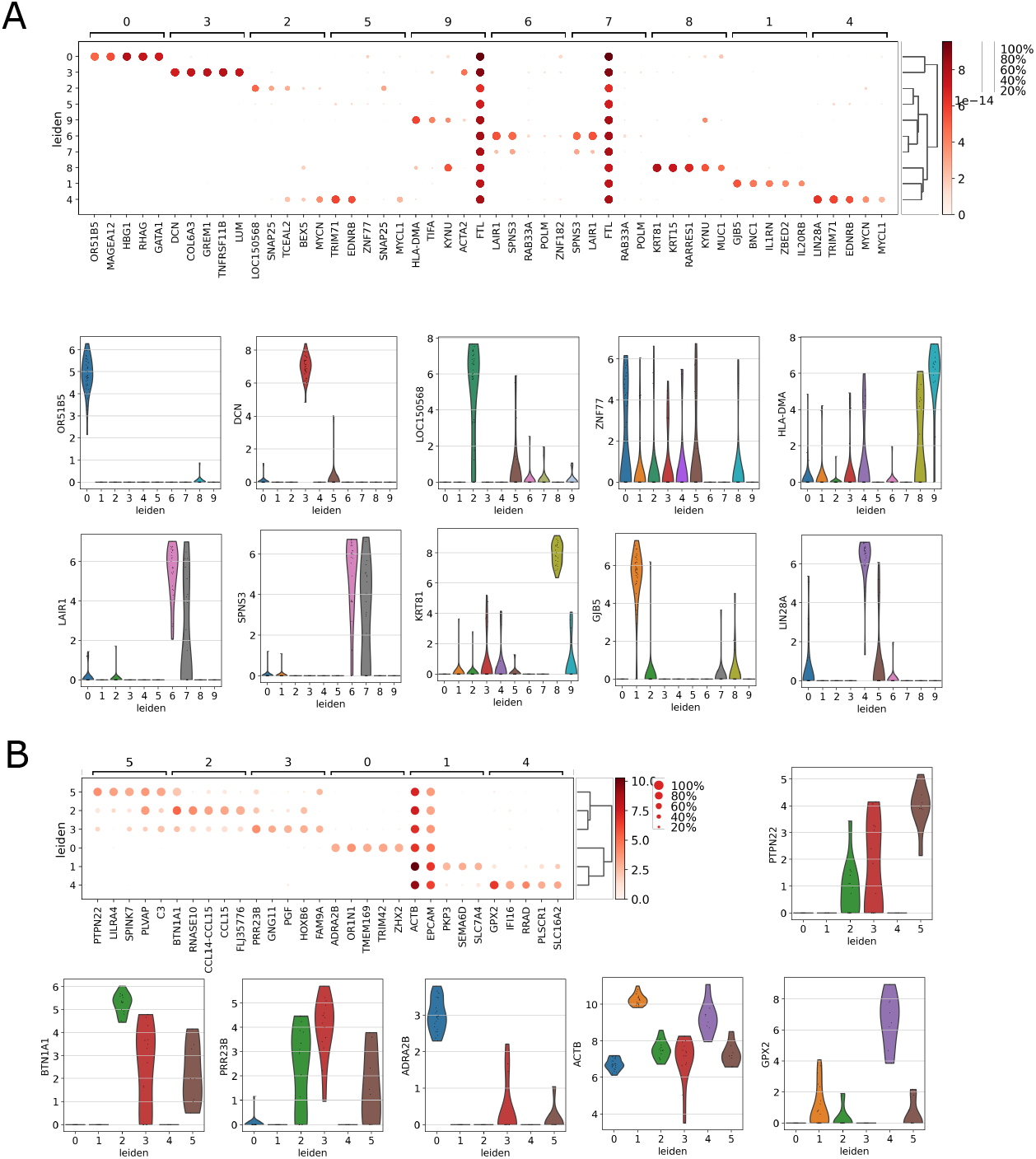
Figure shows marker analysis for Pollen dataset (panel-A), and Yan dataset (panel-B). The average expression values of the top five DE genes are shown in heatmap of panel-A, and -B. The violin plots of the expression profiles of those top DE genes within each cluster are shown in panel-A and -B.

### A case study on ultra large scRNA-seq data

*RgCop* is applied on large PBMC68k dataset with sample size 68000. After preprocessing the data, *RgCop* selects 100 genes which are utilized for further processing. We adopt the conventional steps of Scanpy python package for scRNA-seq clustering. The PBMC cells is clustered by Leiden graph-clustering method [16]. Figure 6, panel-A, -B and -C show the t-SNE visualization of the clustering results for *RgCop*. For comparison purposes, we performed the gene selection step of all other competing methods in PBMC68k data and follow similar steps for clustering.

**Fig 6.**
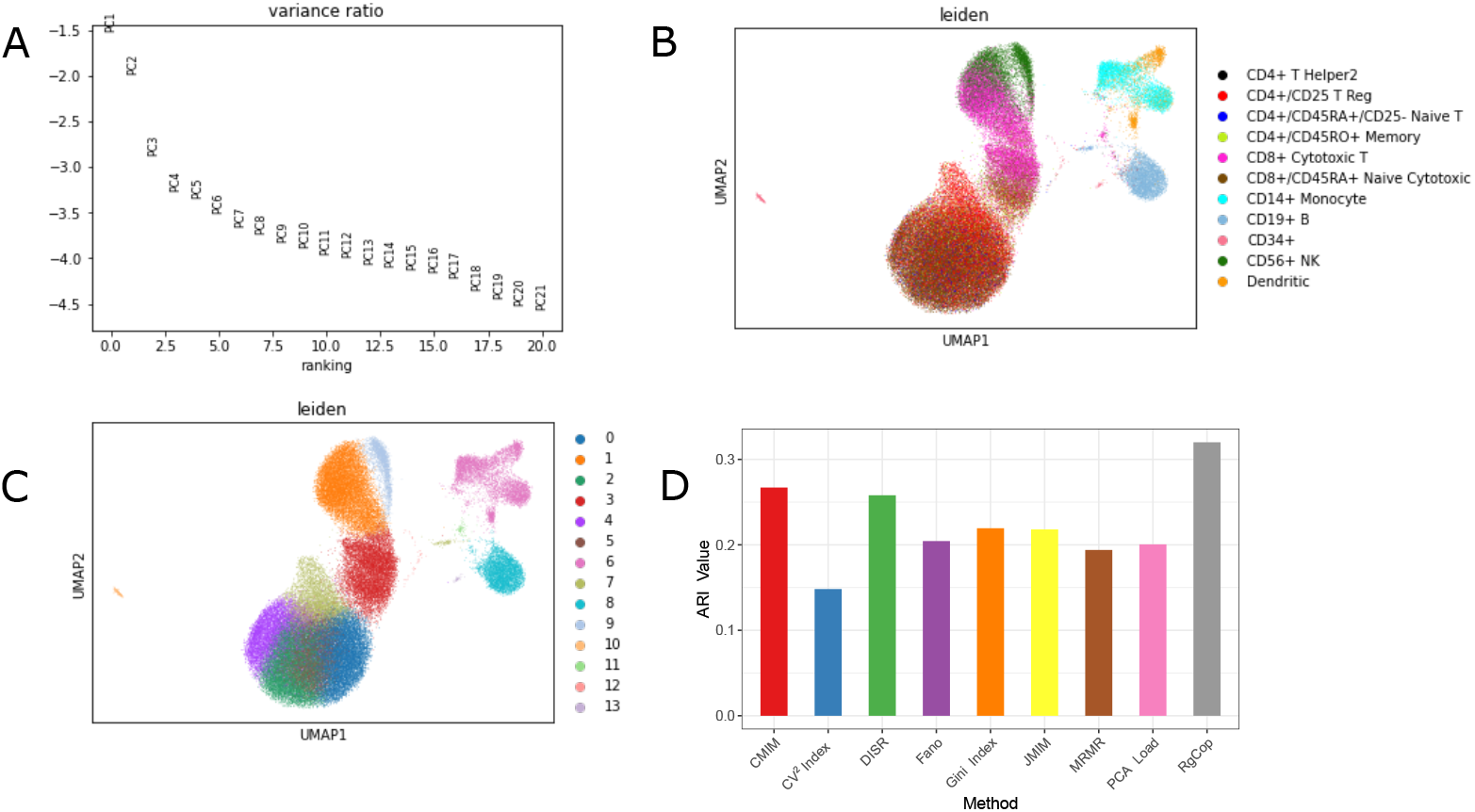
Figure shows the clustering results on large PBMC68k datasets. Panel-A, -B and -C represents the t-sne visualization the clusters obtained from the selected feature sets of *RgCop*. Panel-D shows a comparisons of ARI scores among all the competing methods (all supervised and unsupervised methods).

## Method

### Datasets description

#### Single cell RNA sequence Datasets

The study used the following single-cell RNA sequence datasets: Yan [24], Pollen [25], Muraro [26] and PBMC68k [1] (see table 3). A detailed description of the data is provided in the following text.

- Yan: The dataset consists of a transcriptome of 124 individual cells from a human preimplantation embryo and embryonic stem cell. The 7 unique cell types accommodates labelled 4-cell, 8-cell, zygote, Late blastocyst, and 16-cell.[GEO under accession no. GSE36552; [24]].
- Pollen: Single cell RNA seq pair-end 100 reads from single cell cDNA libraries were quality trimmed using Trim Galore with the flags. It contains 11 cell types. [GEO accession no GSM1832359; [25]]
- Muraro: Single-cell transcriptomics was carried out on live cells from a mixture using an automated version of CEL-seq2 on live, FACS sorted cells. It contains 2126 number of cells. It is a human pancreas cell tissue with 10 cell types. The dataset was downloaded from GEO under accession no GSE85241 [26].
- PBMC68k: The dataset [1], is downloaded from 10x genomics website https://support.10xgenomics.com/single-cell-geneexpression/datasets. The data is sequenced on Illumina NextSeq 500 high output with 20,000 reads per cell.

**Table 3.**
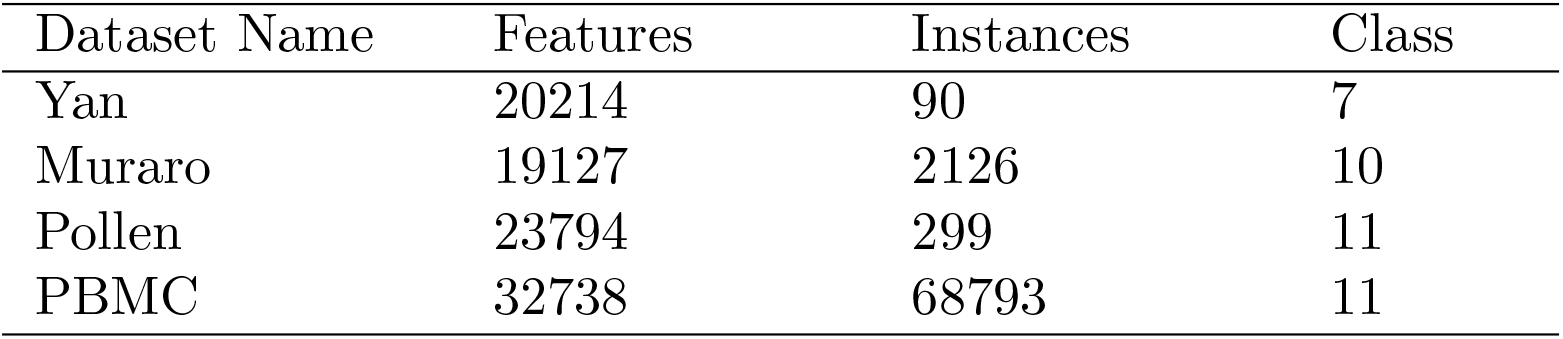
A brief summary of the real scRNA sequence Dataset

### Background theory that supports *RgCop*

#### Short description on Copula

The ‘Copula’ term [27] is originated from a Latin word *copulare*, which joins multivariate distributions to its one dimensional distribution function. The copula is considerably employed in high dimensional datasets to obtain joint distributions using uniform marginal distributions and vice versa. See supplementary for a detailed descriptions of copula and its related measures.

#### Copula correlation measure

Let, *Y* = {*y*_1_*, y*_2_} and *Z* = {*z*_1_*, z*_2_} are two bivariate random variables and their joint and marginal distributions are *H_Y Z_*, *F_Y_* (*y*) and, *F_Z_*(*z*) respectively. Now *H_Y Z_* can be expressed as: *H_Y Z_*(*y, z*) = *C*(*F_Y_*(*y*), *F_Z_*(*z*)), where *C* is a copula function.

Kendall tau(*τ*), the measure of association, [28] can be expressed in terms of concordance and discordance between random variables. Kendall tau is the difference between probability of concordance and discordance of (*y*_1_*, y*_2_) and (*z*_1_*, z*_2_). It can be described as

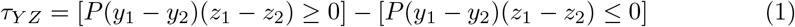

According to Nelson [29] Kendall tau can be expressed using copula function:

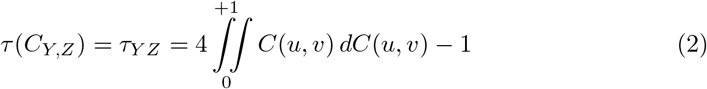

Where, *u ∈ F_Y_* (*y*) and *v ∈ F_Z_* (*z*). *τ*(*C_Y,Z_*) is termed as copula-correlation (*Ccor*) in our study.

#### A note on regularization

Regularization is a type of regression that penalizes the coefficient of redundant feature towards zero [30] (see supplementary for detailed description). The simplest regularization is *l*_1_ norm or Lasso Regression, which adds “absolute value of magnitude” of coefficient as penalty term to the loss function. For any vector 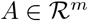, the *l*_1_ norm is 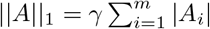, w*^m^*here *γ* is a tuning parameter, controls penalization. For *γ* = 0 regularization effect is none. When *γ* value increases, it starts to penalizes the larger coefficients to zero. However, after a certain value of *γ*, the model starts losing important properties, increasing bias in the model and thus causes under-fitting. We tuned *γ* using eight synthetic Gaussian mixture dataset in this study.

### Gene selection using *RgCop* algorithm

#### Max relevancy

A gene *g_i_* is more relevant to class labels *C_D_* than another gene *g_j_*, if *g_i_* has higher *Ccor* score with *C_D_* than *g_j_*. This is called *Relevancy* test for the genes [31] and is used to select most relevant gene from a gene set. Formally it can be described as: *g_i_* ≺ *g_j_* if the following is true,

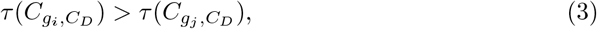

Where *τ*[*C*(*x, y*)] represents copula correlation between two random variable *X* and *Y*. For estimating the copula density we have used empirical copula. The maximum-relevancy method choose the gene (feature) subset among gene set *G* as

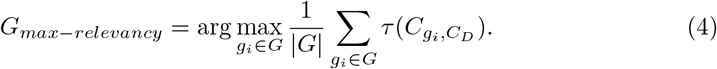

*G_max-relevancy_* may contains genes that are mutually dependent. This is because we only consider the *Ccor* between a gene and class labels, overlooking the mutual dependency among the selected and non-selected genes. This results spurious genes in the selected list.

#### Min Redundancy

Redundancy is a measure that computes the mutual dependence among set random variables. Here we used the same definition of *Ccor* to compute multivariate dependency between selected gene (*g_i_*) and non selected gene sets (*g_s_*). Formally it can be expressed as

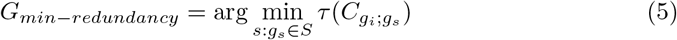

#### Objective function

*RgCop* utilizes a forward selection wrapper approach to select gene iteratively from a gene set. It uses multivariate copula-based dependency instead of the classical information measure. The objective function integrates the relevancy and the redundancy terms defined using the *Ccor*. Mathematically, it can be expressed as follows.

Let us assume genes (*g*_1_*,…, g_i_*) are in the selected list *G_s_*. The next gene *g*_*i*+1_ ∈ (*G* – *G_s_*) in at (*i* + 1) iteration is using the objective function

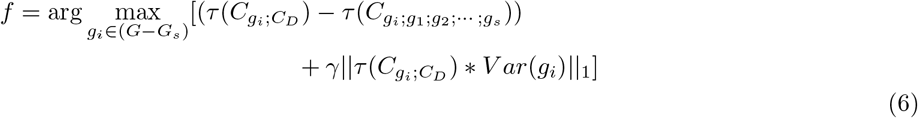

Where, 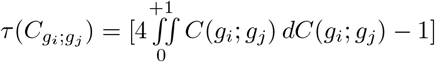 is Kendall tau dependency score of Empirical copula between two genes *g_i_* and *g_j_*. Here, *γ* represents the regularization coefficient. An overview of the *RgCop* algorithm is given in algorithm-1

#### Proof of correctness

Suppose *G_s_* = {*g*_1_, *g*_2_,…, *g_i_, g_i+1_*,…*g_d_*} denotes a subset of genes obtained from a gene set *G* using *RgCop*. Here *g_i_* represents selected gene at iteration *i*. We claim that the set *G_s_* is optimal.

#### Proof

Let us prove this by the method of contradiction. If we assume the claim is not true, then there should exist some another optimal gene set *G′_s_*. Without loss of generality, let us assume *G′_s_* has a maximum number of initial genes (*i* number genes) common with *G_s_*.

Now *G′_s_* can be written as *G′_s_* = {*g*_1_*, g*_2_*,…, g_i_, g_k_,…, g_d_*}. So, *G′_s_* contains {*g*_1_*, g*_2_*,…, g_i_*} from *G_s_*, but not *g*_*i*+1_. Following our assumption *g*_*i*+1_ cannot be included in any of the optimal gene lists (*G′_s_* has maximum *i* number of initial genes overlapped with *G_s_*).

Now we claim that *k* > (*i* + 1). This is because *k* cannot be equal to *i* + 1, otherwise *G′_s_* would have (*i* + 1) genes overlapped with *G_s_*. Similarly, *k* ≰ *i*, because otherwise *G′_s_* will contains redundant genes.

Now by the definition of our objective function (*f*) we can write: *f* (*g_k_*) < *f* (*g*_*i*+1_). So we can substitute *g_k_* with *g*_*i*+1_ in the *G′_s_* list, and the list will be still optimal. This contradicts our assumption that *g*_*i*+1_ cannot be included in any optimal list. This proves our claim.

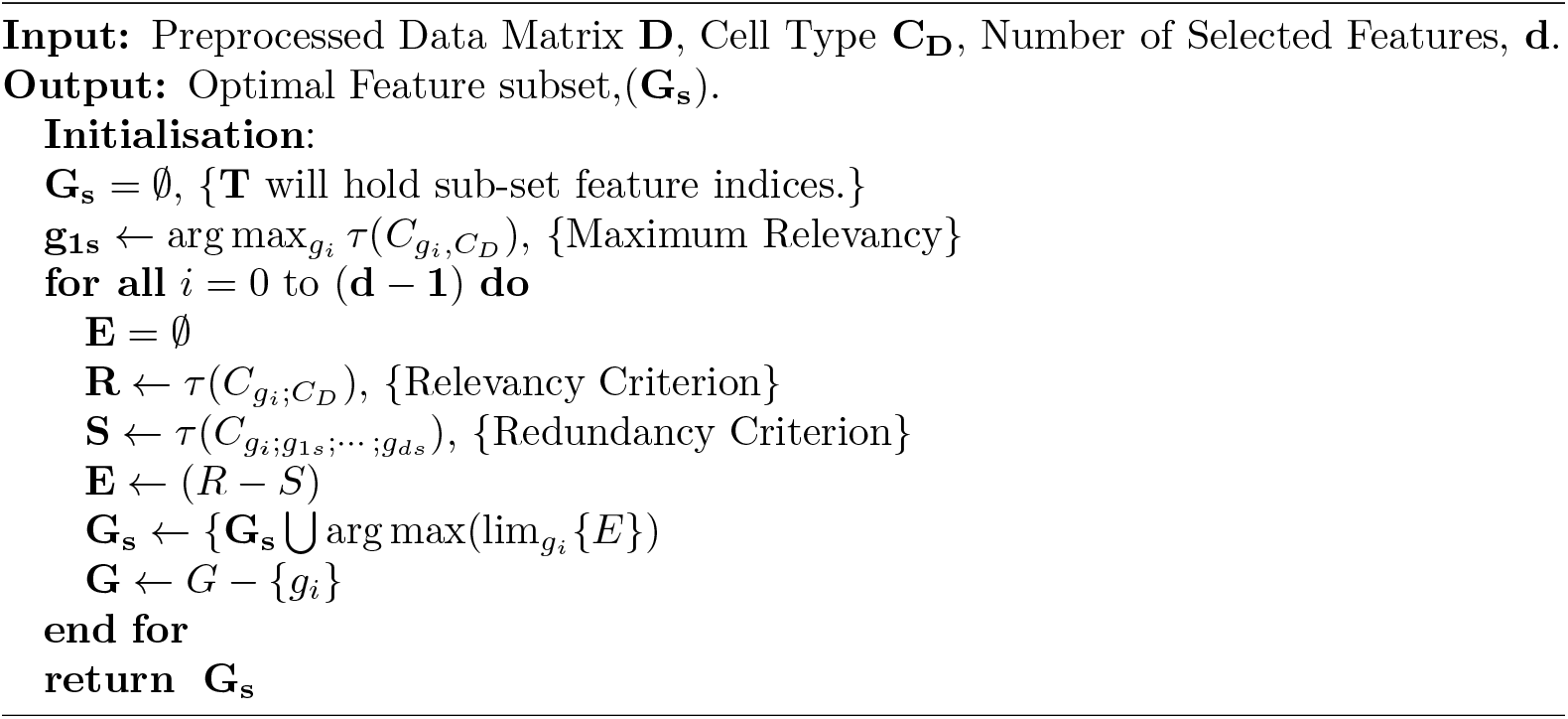

## 2 Conclusions

The selection of informative genes in scRNA-seq data is crucial and an essential step for the downstream analysis. Because of the large feature/gene set of scRNA-seq data, selecting important genes is a challenging task which has an immense effect in clustering and annotation results in the later stage of downstream analysis. The proposed method *RgCop* addressed this task by employing a robust and equitable dependence measure called copula-correlation (*Ccor*). It can accurately measure relevancy and redundancy simultaneously between two sets of gene. *RgCop* also add simple *l*_1_ regularization technique with its objective function to control the large coefficients of relevancy terms. Realistic simulations confirm the utmost accuracy of the *RgCop* in overlapped and non-overlapped Gaussian mixture data. We also demonstrated that *RgCop* results high accuracy both in clustering and classification performance with the selected genes from real-life scRNA-seq datasets. The identified marker genes can also be able to dissect cell clusters, suggesting the inclusions of marker genes within the selected sets.

*RgCop* introduces a stable feature/gene selection which is evaluated by applying it in noisy data. By virtue of the important *scale invariant property* of copula, the selected features are invariant under any transformation of data due to the most common technical noise present in the scRNA-seq experiment. The range of tuning parameter (regularization coefficient (*λ*)) is determined using *RgCop* on synthetic Gaussian mixture dataset. *RgCop* also produces accurate clustering/classification results on four sc-RNA seq datasets. The results are validated using ARI score/classification accuracy. The stability of *RgCop* is evaluated by applying it in noisy data and matching the resulting feature set with the original one. This was performed multiple times with varying number of selected features. The resulting ARI scores utilize a minimum deviation suggesting a robust and stable approach for feature selection.

It can be noted that although *RgCop* primarily detect variable genes from scRNA-seq data, we extended the process by employing a clustering/classification technique with it to annotate unknown cells. The efficacy of *RgCop* is demonstrated by applying it to cluster/group unknown cells with the selected genes/features. A precise annotation of cell clusters also illustrates the applicability of *RgCop* to select the most variable genes in the early stage. The most general classifiers trained with the selected features can accurately predict the cell types of an unknown sample. Several genes are highlighted having a high expression level within clusters, which are acting as markers.

Taken together, the proposed method *RgCop* not only outperforms in informative gene selection but also can able to annotate unknown cells/cell-clusters in scRNA-seq data. It can be observed from the results that *RgCop* leads both in the domain of robust gene/feature selection and type annotation of unknown cell in large scRNA-seq. The results prove that *RgCop* may be treated as an important tool for computational biologist to investigate the primary steps of downstream analysis of scRNA-seq data.

## Notes

### Competing Interest Statement

The authors have declared no competing interest.

## References

1. Zheng GX, Terry JM, Belgrader P, Ryvkin P, Bent ZW, Wilson R, et al. Massively parallel digital transcriptional profiling of single cells. Nature communications. 2017;8:14049.

2. Lall S, Ghosh A, Ray S, Bandyopadhyay S. sc-REnF: An Entropy Guided Robust Feature Selection for Single-Cell RNA-seq Data. bioRxiv. 2021;.

3. Ray S, Bandyopadhyay S, et al. Generating realistic cell samples for gene selection in scRNA-seq data: A novel generative framework. bioRxiv. 2021;.

4. Kiselev VY, Kirschner K, Schaub MT, Andrews T, Yiu A, Chandra T, et al. SC3: consensus clustering of single-cell RNA-seq data. Nature methods. 2017;14(5):483.

5. Ji Z, Ji H. TSCAN: Pseudo-time reconstruction and evaluation in single-cell RNA-seq analysis. Nucleic acids research. 2016;44(13):e117–e117.

6. Plass M, Solana J, Wolf FA, Ayoub S, Misios A, Glažar P, et al. Cell type atlas and lineage tree of a whole complex animal by single-cell transcriptomics. Science. 2018;360(6391).

7. Fincher CT, Wurtzel O, de Hoog T, Kravarik KM, Reddien PW. Cell type transcriptome atlas for the planarian Schmidtea mediterranea. Science. 2018;360(6391).

8. Ray S, Schonhuth A. MarkerCapsule: Explainable Single Cell Typing using Capsule Networks. bioRxiv. 2020;.

9. Luecken MD, Theis FJ. Current best practices in single-cell RNA-seq analysis: a tutorial. Molecular systems biology. 2019;15(6):e8746.

10. Butler A, Hoffman P, Smibert P, Papalexi E, Satija R. Integrating single-cell transcriptomic data across different conditions, technologies, and species. Nature biotechnology. 2018;36(5):411–420.

11. Kim JM, Jung YS, Sungur EA, Han KH, Park C, Sohn I. A copula method for modeling directional dependence of genes. BMC bioinformatics. 2008;9(1):225.

12. Ray S, Lall S, Bandyopadhyay S. CODC: a Copula-based model to identify differential coexpression. NPJ systems biology and applications. 2020;6(1):1–13.

13. Kasa SR, Bhattacharya S, Rajan V. Gaussian mixture copulas for high-dimensional clustering and dependency-based subtyping. Bioinformatics. 2020;36(2):621–628.

14. Yip SH, Wang P, Kocher JPA, Sham PC, Wang J. Linnorm: improved statistical analysis for single cell RNA-seq expression data. Nucleic acids research. 2017;45(22):e179–e179.

15. Wolf FA, Angerer P, Theis FJ. SCANPY: large-scale single-cell gene expression data analysis. Genome biology. 2018;19(1):15.

16. Traag VA, Waltman L, van Eck NJ. From Louvain to Leiden: guaranteeing well-connected communities. Scientific reports. 2019;9(1):1–12.

17. Jiang L, Chen H, Pinello L, Yuan GC. GiniClust: detecting rare cell types from single-cell gene expression data with Gini index. Genome biology. 2016;17(1):144.

18. Macosko EZ, Basu A, Satija R, Nemesh J, Shekhar K, Goldman M, et al. Highly parallel genome-wide expression profiling of individual cells using nanoliter droplets. Cell. 2015;161(5):1202–1214.

19. Grün D, Kester L, Van Oudenaarden A. Validation of noise models for single-cell transcriptomics. Nature methods. 2014;11(6):637.

20. Fleuret F. Fast binary feature selection with conditional mutual information. Journal of Machine Learning Research. 2004;5(Nov):1531–1555.

21. Bennasar M, Hicks Y, Setchi R. Feature selection using joint mutual information maximisation. Expert Systems with Applications. 2015;42(22):8520–8532.

22. Meyer PE, Bontempi G. On the use of variable complementarity for feature selection in cancer classification. In: Workshops on Applications of Evolutionary Computation. Springer; 2006. p. 91–102.

23. Peng H, Long F, Ding C. Feature selection based on mutual information criteria of max-dependency, max-relevance, and min-redundancy. IEEE Transactions on pattern analysis and machine intelligence. 2005;27(8):1226–1238.

24. Yan L, Yang M, Guo H, Yang L, Wu J, Li R, et al. Single-cell RNA-Seq profiling of human preimplantation embryos and embryonic stem cells. Nature structural & molecular biology. 2013;20(9):1131.

25. Pollen AA, Nowakowski TJ, Shuga J, Wang X, Leyrat AA, Lui JH, et al. Low-coverage single-cell mRNA sequencing reveals cellular heterogeneity and activated signaling pathways in developing cerebral cortex. Nature biotechnology. 2014;32(10):1053.

26. Muraro MJ, Dharmadhikari G, Grün D, Groen N, Dielen T, Jansen E, et al. A single-cell transcriptome atlas of the human pancreas. Cell systems. 2016;3(4):385–394.

27. Nelsen RB. An introduction to copulas. Springer Science & Business Media; 2007. 429

28. Kruskal WH. Ordinal measures of association. Journal of the American Statistical Association. 1958;53(284):814–861.

29. Nelsen RB. Properties and applications of copulas: A brief survey. In: Proceedings of the First Brazilian Conference on Statistical Modeling in Insurance and Finance,(Dhaene, J., Kolev, N., Morettin, PA (Eds.)), University Press USP: Sao Paulo; 2003. p. 10–28.

30. Xing E, Jordan M, Karp I, Richard M. Feature selection for high-dimensional genomic microarray data. In: ICML. vol. 1; 2001. p. 601–608.

31. Brown G, Pocock A, Zhao MJ, Luján M. Conditional likelihood maximisation: a unifying framework for information theoretic feature selection. Journal of machine learning research. 2012;13(Jan):27–66.

